# The first near-complete assembly of the hexaploid bread wheat genome, *Triticum aestivum*

**DOI:** 10.1101/159111

**Authors:** Aleksey V. Zimin, Daniela Puiu, Richard Hall, Sarah Kingan, Bernardo J. Clavijo, Steven L. Salzberg

**Affiliations:** Center for Computational Biology, McKusick-Nathans Institute of Genetic Medicine, Johns Hopkins University School of Medicine, Baltimore, MD; Institute for Physical Sciences and Technology, University of Maryland, College Park, MD; Pacific Biosciences, Menlo Park, CA; Earlham Institute, Norwich NR4 7UZ, United Kingdom; Departments of Biomedical Engineering, Computer Science, and Biostatistics, Johns Hopkins University, Baltimore, MD

## Abstract

Common bread wheat, *Triticum aestivum*, has one of the most complex genomes known to science, with 6 copies of each chromosome, enormous numbers of near-identical sequences scattered throughout, and an overall size of more than 15 billion bases. Multiple past attempts to assemble the genome have failed. Here we report the first successful assembly of *T. aestivum*, using deep sequencing coverage from a combination of short Illumina reads and very long Pacific Biosciences reads. The final assembly contains 15,344,693,583 bases and has a weighted average (N50) contig size of of 232,659 bases. This represents by far the most complete and contiguous assembly of the wheat genome to date, providing a strong foundation for future genetic studies of this important food crop. We also report how we used the recently published genome of *Aegilops tauschii*, the diploid ancestor of the wheat D genome, to identify 4,179,762,575 bp of *T. aestivum* that correspond to its D genome components.

## Introduction

For many years, the hexaploid (AABBDD) bread wheat genome, *Triticum aestivum*, has resisted efforts to sequence and assemble it. The first effort to sequence the genome, published in 2012 [1], used an earlier generation of sequencing technology and only assembled 5.42 billion bases (Gbp), approximately one-third of the genome. In a second attempt two years later, an international consortium published the results of a systematic effort to sequence the genome one chromosome at a time, using deep coverage in 100-bp Illumina reads [2]. That effort, although more successful than the previous one, yielded only 10.2 billion bases of sequence, approximately two-thirds of the genome. The contiguity of this assembly was quite poor, with the 10.2 billion bases divided amongst hundreds of thousands of contigs, and with N50 sizes ranging from 1.7 to 8.9 kilobases (Kb) for the different chromosome arms. In 2017, a third assembly of wheat was published, estimated to represent 78% of the genome [3]. This assembly contained 12.7 billion bases of sequence, but it too was highly fragmented, containing over 2.7 million contigs with an N50 contig size of 9,731 bp.

The wheat genome’s complexity, and the challenge it presents for genome assembly, stems not only from its large size (five times the size of the human genome), but also from its very high proportion of relatively long, near-identical repeats, most of them due to transposable elements [4]. Because these repeats are much longer than the length of Illumina reads, efforts to assemble the genome using Illumina data have been unable to resolve these repeats. Another major challenge in assembling the wheat genome is that it is hexaploid, and the three component genomes–wheat A, B, and D, each comprising seven chromosomes–share many regions of high similarity. Genome assembly programs are thus faced with a doubly complex problem: first that the genome is unusually repetitive, and second that each chromosome exists in six copies with varying degrees of intra- and inter-chromosome similarity.

The most effective way to resolve repeats is to generate individual reads that contain them. If a single read is longer than a repeat, and if both ends of the read contain unique sequences, then genome assemblers can unambiguously place the repeat in the correct location. Without such reads, every long repeat creates a breakpoint in the assembly. Recent advances in sequencing, particularly the long read, single-molecule sequencing technologies from Pacific Biosciences (PacBio) and Oxford Nanopore, can produce reads in excess of 10,000 bp, although with a high error rate. By combining these very long reads with highly accurate shorter reads, we have been able to produce an assembly of the wheat genome that is dramatically better than any previous attempt. Ours is the first assembly that contains essentially the entire length of the genome, with more than 15.3 billion bases, and its contiguity is more than *ten times* better than the partial assemblies published in the past.

## Results

To create the wheat genome assembly, we generated two extremely large primary data sets. The first data set consisted of 7.06 billion Illumina reads containing approximately 1 trillion bases of DNA. The Illumina reads were 150-bp, paired reads from short DNA fragments, averaging 400 bp in length. Using an estimated genome size of 15.3 Gbp, this represented 65-fold coverage of the genome. The second data set used Pacific Biosciences single-molecule (SMRT) technology to generate 55.5 million reads with an average read length just under 10,000 bp, containing a total of 545 billion bases of DNA, representing 36-fold coverage of the genome. All reads were generated from the Chinese spring variety (CS42, accession Dv418) of *T. aestivum*, the same variety as used in earlier attempts to sequence the genome.

### MaSuRCA assembly

To create the initial assembly, Triticum 1.0, we ran the MaSuRCA assembler (v. 3.2.1) on the full data set of Illumina and PacBio reads. The first major step was the creation of super-reads [5] from the Illumina reads. Super-reads are highly accurate and longer than the original reads, and because they are much fewer in number, they provide a means to greatly compress the original data. This step generated 95.7 million super-reads with a total length of 31 Gb, a mean size of 324 bp and an N50 size of 474 bp (i.e., half of the total super-read sequence was contained in super-reads of 474 bp or longer). The super-reads provided a 32-fold compression of the original Illumina data.

Next we created *mega-reads* by using the super-reads to tile the PacBio reads, effectively replacing most PacBio reads (which have an average error rate of ~15%) with much more accurate sequences [6]. Most PacBio reads were converted into a single mega-read, but in some cases a given PacBio read yielded two or more (shorter) mega-reads. In total we created 57,020,767 mega-reads with a mean length of 4,876 bp and an N50 length of 8,427 bp. The total length of the mega-reads was 278 Gb, representing about 18X genome coverage. As part of this step, we also created synthetic mate pairs; these link together two mega-reads when the pair of mega-reads originates from a single PacBio read. We generated these pairs by extracting 400 bp from opposite ends of each pair of consecutive mega-reads corresponding to a given PacBio read. This resulted in 23.45 million pairs of 400 bp reads, totalling 18.75 Gb.

Construction of super-reads and mega-reads required approximately 100,000 CPU hours, of which 95% was spent in the mega-reads step. By using large multi-core computers to run these steps in parallel, these steps took 1.5 months of elapsed (wall clock) time. The peak memory (RAM) usage was 1.2 terabytes.

We then assembled the mega-reads and the synthetic pairs using the Celera Assembler [7] (v8.3), which was modified to work with our parallel job scheduling system. The CA assembly process required many iterations of the overlapping, error correction, and contig construction steps, and it was extremely time consuming, even with the many optimizations that have been incorporated in this assembler in recent releases. The total CPU time was ~470,000 CPU hours (53.7 *years*), which was only made feasible by running it on a grid with thousands of jobs running in parallel for some of the major steps. The total elapsed time was just over 5 months. When combined with the earlier steps, the entire assembly process took 6.5 months. The resulting assembly, labelled Triticum 1.0, contained 17.046 Gb in 829,839 contigs, with an N50 contig size of 76,267 bp and an N50 scaffold size of 101,195 bp (**Table 1**).

Next, in order to detect and remove redundant regions of the assembly, we aligned the assembly against itself using the nucmer program from the MUMmer package [8]. We identified and excluded scaffolds that were completely contained in and ≥96% identical to other scaffolds. After this de-duplication procedure, the reduced assembly, Triticum 2.0, contained 14.40 Gbp in 375,328 contigs with an N50 contig size of 75,599 bp, with scaffolds spanning 14.45 Gbp and an N50 scaffold size of 100,805 bp (**Table 1**).

**Table 1.** Assembly statistics for each of the assemblies of *Triticum aestivum* constructed as described in the text. To enable fair comparisons, all N50 sizes are computed using an estimated genome size of 15.34 Gb.

|  |  |  |  |  |  |
| --- | --- | --- | --- | --- | --- |
| Assembly | Element type | Number | Total size (bp) | Average size (bp) | N50 size (bp) |
| Triticum 1.0 | contigs | 829,839 | 17,045,571,778 | 20,541 | 76,267 |
|  | scaffolds>2Kb | 576,137 | 16,889,295,941 | 29,314 | 101,195 |
| Triticum 2.0 | contigs | 375,328 | 14,395,027,822 | 38,353 | 75,599 |
|  | scaffolds>2Kb | 252,501 | 14,412,484,332 | 57,078 | 100,805 |
| FALCON Trit 1.0 | contigs | 97,809 | 12,939,100,857 | 132,289 | 215,314 |
| Triticum 3.0 | contigs | 279,439 | 15,343,711,528 | 54,908 | 232,613 |
| Triticum 3.1 | contigs | 279,439 | 15,344,693,583 | 54,912 | 232,659 |

### FALCON assembly

Independently of the MaSuRCA assembly, we assembled the PacBio data alone using the FALCON assembler [9], followed by polishing with the Arrow program, which substantially improves the consensus accuracy. FALCON implements a hierarchical assembly approach; the initial step is to error correct long reads by aligning all reads to a subset of the longest reads. Given the relatively low raw read coverage (36X), we used a long-read cutoff of 1 Kb, generating 11X coverage of error-corrected reads with an N50 size of 16 Kb. Error correction and assembly of the corrected reads was completed using ~150,000 CPU hours, which took ~3 weeks on a 16-node cluster. The contigs output from FALCON require further polishing, which involves realignment of raw reads and calculation of a new consensus [10]. For the polishing step, we used Pacbio’s resequencing pipeline from the SMRT Analysis package (https://github.com/PacificBiosciences/SMRT-Link) after first splitting the assembled contigs into <4 Gbp chunks (a limit of the aligner). Polishing required an additional ~160,000 CPU hours, for a total of 310,000 CPU hours and 6 weeks elapsed (wall clock) time.

These steps produced an assembly, designated FALCON Trit 1.0, containing 12.94 Gbp in 97,809 contigs with a mean size of 132,289 and an N50 size of 215,314 bp (**Table 1**).

### Merged assembly

The contigs from the FALCON assembly were larger than those from the MaSuRCA assembly; however, the total size of the assembly was 1.5 Gbp smaller. To capture the advantages of both assemblies, we merged them as follows. We aligned the contigs (not scaffolds) from the two assemblies using MUMmer 4.0 [8] and extracted all pairwise best matches. We then merged each pair of FALCON contigs when they overlapped a single Triticum 2.0 contig by at least 5000 bp, with Triticum 2.0 sequence filling the gap (see **Figure 1**).

**Figure 1.**
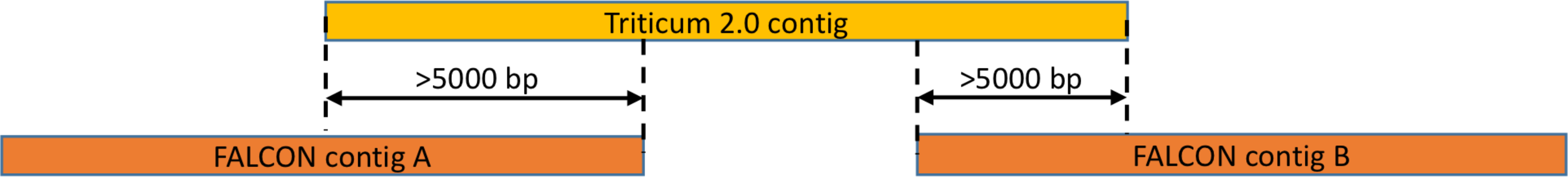
Illustration of the merging process for the Triticum 2.0 and FALCON Trit 1.0 assemblies. If two contigs A and B from the FALCON assembly overlapped a Triticum 2.0 contig by at least 5000 bp, then A and B were merged together, using the Triticum 2.0 contig to fill the gap.

After merging and extending the FALCON contigs, we then identified all MaSuRCA scaffolds that were not contained in the longer FALCON contigs, and added these to the new assembly. The resulting merged assembly, Triticum 3.0, contains 15,343,750,409 bp in 279,529 contigs, with a contig N50 size of 232,613 bp (**Table 1**). The longest contig is 4,510,883 bp.

### Genome complexity

As described above, previous attempts to assemble the hexaploid wheat genome were stymied because of its exceptionally high repetitiveness, but until now we had no reliable way to quantify how repetitive the genome truly is. To answer this question with a precise metric, we computed the k-mer uniqueness ratio, a metric defined earlier as a way to capture repetitiveness that reflects the difficulty of assembly [11]. This ratio is defined as the percentage of a genome that is covered by unique sequences of length *k* or longer. If, for example, 90% of a genome is comprised of unique 50-mers, then one might expect that 90% of that genome could be assembled using accurate (low-error-rate) reads that were longer than 50 bp.

With the Triticum 3.0 assembly in hand, we computed the k-mer uniqueness ratio for wheat and compared it to several other plant and animal genomes, as shown in **Figure 2**. As the figure illustrates, for any value of k, a much smaller percentage of the wheat genome is covered by unique k-mers than other plant or animal genomes, with the exception of *Ae. tauschii*, which as expected (because it is near-identical to the D genome of hexaploid *T. aestivum*) is only slightly less repetitive. For example, only 44% of the 64-mers in the wheat genome are unique, as contrasted with 90% of the 64-mers in cow and 81% of the 64-mers in rice. This analysis demonstrates that in order to obtain an assembly covering most of the wheat genome, particularly if the algorithm relies on de Bruijn graphs, much longer reads will be required. Our sequencing strategy, by using deep coverage in very long PacBio reads coupled with highly accurate Illumina reads, was able to produce the long, accurate reads required to assemble this very complex genome.

**Figure 2.**
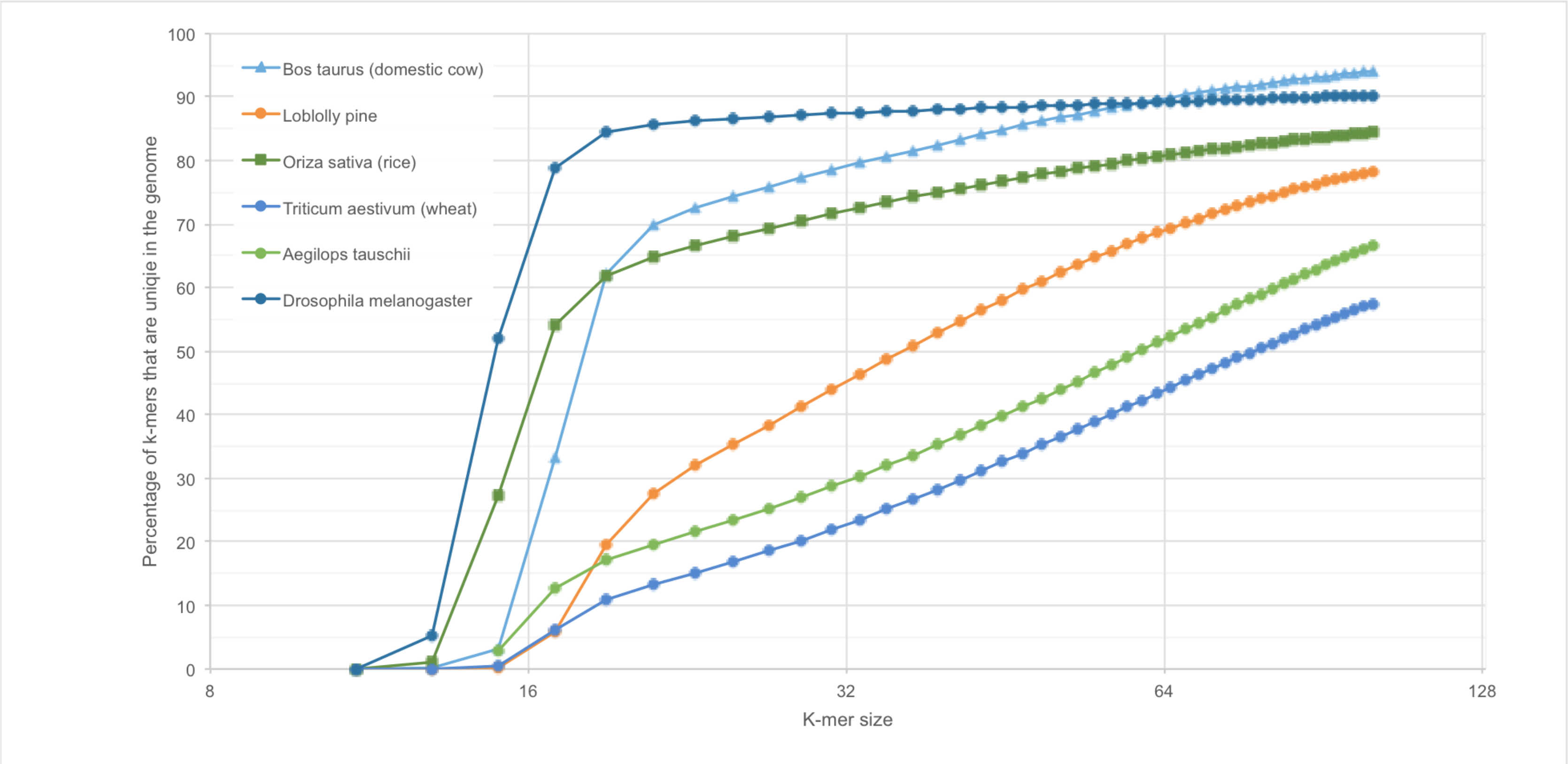
K-mer uniqueness ratios for the wheat genome (*Triticum aestivum*) compared to the cow, fruit fly, rice, loblolly pine, and *Ae. tauschii* genomes. The plot shows the percentage of each genome that is covered (y-axis) by unique sequences of length k, for various values of k (x-axis).

### Identifying the wheat D genome

*T. aestivum* is a hexaploid plant with three diploid ancestors, one of which is *Aegilops tauschii*, commonly known as goat grass. *Ae. tauschii* itself is a highly repetitive genome that has resisted attempts at assembly, but we recently published a highly contiguous draft assembly (Aet_MR 1.0) using a similar strategy to the one used for wheat, a combination of PacBio and Illumina sequences [6]. *T. aestivum*’s hexaploid composition is typically represented as AABBDD, where the D genome was contributed by an ancestor of *Ae. tauschii*. The hexaploidization event occurred very recently, approximately 8,000 years ago, when *Ae. tauschii* spontaneously hybridized with a tetraploid wheat species, *Triticum turgidum* [12].

Because this event was so recent, the wheat D genome and *Ae. tauschii* are highly similar, much closer to one another than the D genome is to either the A or B genomes. We used this similarity to identify the D genome components of our assembly by aligning the *Ae. tauschii* contigs in Aet_MR 1.0 to Triticum 3.0. We used the nucmer program [8] to identify all alignments representing best matches between Triticum 3.0 and Aet_MR 1.0 with a minimum identity of 97%. The vast majority of the two genomes are >99% identical, making this filtering process relatively straightforward.

After filtering, we identified 50,101 contigs with a total length of 4,179,762,575 bp from Triticum 3.0 that aligned to *Ae. tauschii*. We separated these D genome contigs from Triticum 3.0 and provided them as the first release of the wheat D genome, which we have named TriticumD 1.0. The N50 size of these contigs is 224,953 bp, using a genome size estimate of 4.18 Gb for wheat D. The total size of 4.18 Gb corresponds closely to the 4.33 Gb in the recently published *Ae. tauschii* (Aet_MR 1.0) assembly [6].

We also ran the alignments in the other direction, aligning all of Aet_MR 1.0 to TriticumD 1.0, and found that 99.8% of the *Ae. tauschii* assembly matches TriticumD; only 8.96 Mb failed to align. The overall mapping is complex; although most of the *Ae. tauschii* and wheat D genomes align in a 1-to-1 mapping, many scaffolds align in a many-to-one or one-to-many arrangement. Thus the additional 150 Mb in *Ae. tauschii* appears to be due to gain/loss of repeats rather than loss of unique sequence from wheat D.

***Assembly quality***. Assessing the quality of an assembly is challenging, especially when the previous assemblies are so much more fragmented, as they are in the case of *T. aestivum*. However, the very high-fidelity alignments between Triticum 3.0 and the published *Ae. tauschii* genome, at over 99% identity, provide strong support for its accuracy. We found no large-scale structural disagreements between the assemblies, other than the many-to-one mappings for some of the scaffolds. These could indicate that one assembly has over-collapsed a repeat, but they could also indicate a true polymorphism; we do not have sufficient data to distinguish these possibilities. The fact that 99.8% of *Ae. tauschii* aligns to Triticum 3.0 supports the hypothesis that the assembly is largely complete as well.

### Re-polishing to create Triticum 3.1

Finally, we used an independent set of Illumina 250-bp reads from an earlier study [3] to measure the quality of the consensus sequence. We used the KAT program [13] to count all 31-mers in each assembly and compare these counts to the 31-mers in the read data. Because the read data here represented 30-fold coverage of the genome, 31-mers that occur approximately 30 times should represent unique sequences; i.e., they are expected to occur exactly once in the assembly.

The KAT analysis revealed that the FALCON Trit 1.0 assembly was missing a relatively large number of 31-mers that occurred in the reads (**Figure 3**), while the Triticum 2.0 assembly was missing far fewer of these 31-mers. The Triticum 3.0 assembly, which used the polished FALCON assembly for most of its consensus sequence, was also missing many 31-mers. The mostly likely explanation for this effect is that the polishing process over-corrected by replacing some 31-mers with near-identical ones. This would have the effect of creating an excess of 31-mers that occur exactly twice in the assembly, although their coverage indicated that they should occur once. The KAT analysis confirmed this expectation (data not shown).

**Figure 3.**
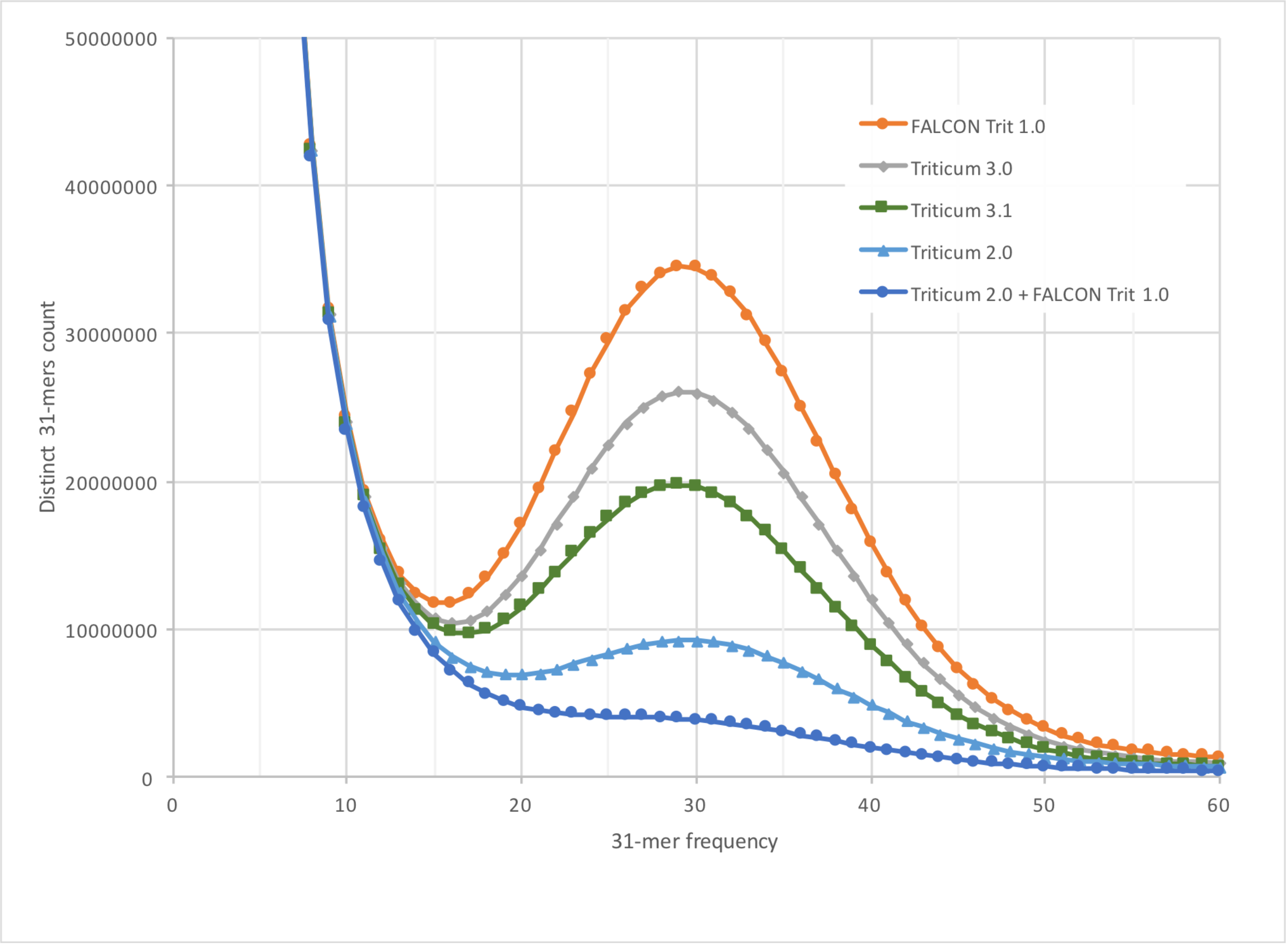
Missing 31-mers in the different assemblies of *Triticum aestivum*. Using the Illumina read data from a previously published assembly of the same genome, we counted all 31-mers in the reads, and then plotted how many of these reads are missing from each assembly. The x-axis shows how often the k-mers occur in the reads. The y-axis shows how many distinct k-mers are missing from each assembly. The FALCON Trit 1.0 assembly had the largest number of missing k-mers, while Triticum 2.0 had the fewest.

We also observed that Triticum 2.0, which used MaSuRCA to create the consensus from Illumina reads, had far fewer missing 31-mers. We therefore re-polished Triticum 3.0 by aligning it to Triticum 2.0, extracting the mutual best matches, and then using the 2.0 sequence as the final consensus. This allowed us to re-polish approximately 11.6 Gbp of the assembly. The resulting assembly, Triticum 3.1, has exactly the same contigs and scaffolds (**Table 1**) but has an improved overall consensus, containing more of the true 31-mers (**Figure 3**). Because of changes in the consensus sequence, the 3.1 assembly is very slightly larger as well. To evaluate the possibility of further improvements, we analysed the 31-mer spectra of both FALCON Trit 1.0 and Triticum 2.0 as a single sequence set. We found that this almost completely eliminated the missing 31-mers (**Figure 3**), illustrating that further improvements in the consensus are possible and are planned for future assembly releases.

## Discussion

In 2004, an international consortium determined that whole-genome shotgun (WGS) sequencing of hexaploid wheat was simply too difficult, "mainly because of the large size and highly repetitive nature of the wheat genome" [14]. The consortium instead determined that the chromosome-by-chromosome approach would be more effective. This strategy, which was far slower and more costly than WGS sequencing, in the end produced a genome assembly that was highly fragmented and that contained only 10.2 Gb [2].

The assembly described here is the first to successfully reconstruct essentially all of the hexaploid wheat genome, *Triticum aestivum*, and to produce relatively large contiguous sequences. The final assembly contains 15,344,693,583 bp with an N50 contig size of 232,659 bp. The previous chromosome-based assembly was not only much smaller overall, but it had average contig sizes approximately 50 times smaller [2]. A recent whole-genome assembly based on deep Illumina sequencing contained 2,726,911 contigs spanning 12,658,314,504 bp and had a contig N50 size of 9731 bp [3]. Compared to Triticum 3.0, that assembly is 2.69 Gb smaller, and its contigs are 24 times smaller. (Note that in order to provide a fair comparison, all N50 sizes reported here are based on the same 15.34 Gb total genome size.)

Why did previous attempts to assemble *T. aestivum* produce a result that was billions of nucleotides shorter than the true genome size? The most likely explanation is that the repetitive sequences, which cover some 90% of the genome [4, 14], are so similar to one another that genome assembly programs cannot avoid collapsing them together. This is a well-known problem for genome assembly, particularly when using the short reads produced by next-generation sequencing technologies. If the differences between repeats occur at a lower rate than sequencing errors, then assemblers cannot distinguish them. The result is an assembly that is both highly fragmented and too short. The same phenomenon can be seen in attempts to assemble *Ae. tauschii*. from short reads. An assembly of that genome using Illumina and 454 sequencing data, contained only 2.69 Gb and had an N50 contig size of just 2.1 Kb [12]. A hybrid assembly using both Illumina and PacBio data, reported by our group early in 2017, produced an assembly of 4.33 Gb, closely matching the estimated genome size, with a contig N50 size of 487 Kb [6].

The key factor in producing a true draft assembly for this exceptionally repetitive genome was the use of very long reads, averaging just under 10,000 bp each, which were required to span the long, ubiquitous repeats in the wheat genome. Deep coverage in these reads (36X, or 545 Gb of raw sequence) coupled with even deeper coverage (65X) in low-error-rate short reads, allowed us to produce a highly accurate and highly contiguous consensus assembly. The massive data set, over 1.5 trillion bases, also required an unprecedented amount of computing power to assemble, and its completion would not have been possible without the availability of very large parallel computing grids. All together, the various assembly steps took 880,000 CPU hours, or just over 100 CPU years. An important technical note is that the computational cost was not simply a function of genome size, but more critically a function of its repetitiveness. The presence of large numbers of unusually long exact and near-exact repeats (Figure 2) means that all of these sequences overlap one another, leading to a quadratic increase in the number of sequence alignments that an assembler must consider.

Finally, by aligning this assembly to the draft genome of *Aegilops tauschii*, the progenitor of the wheat D genome, we were able to cleanly separate the D genome component from the A and B genomes of hexaploid wheat, which is reported here for the first time. This separation was possible because Ae. tauschii is much closer to wheat D, having diverged approximately 8,000 years ago [14], than either genome is to wheat A or B.

The wheat genome presented here provides, for the first time, a near-complete substrate for future studies of this important food crop. Previous efforts to annotate the genome have been hampered by the absence of a large proportion of the genome itself, making inferences about missing genes or gene families difficult, and also by the highly fragmented nature of previous assemblies, which had average contig sizes under 10 Kb. With over half of the genome now contained in contigs longer than 232 Kb, the Triticum 3.0 assembly will contain many more genes within single contigs, greatly aiding future efforts, which are already under way, to study its gene content, evolution, and relationship to other plant species.

**Availability of data**. The Triticum project data have been deposited at the National Center for Biotechnology Information (NCBI) under BioProject PRJNA392179. The assembly has been deposited at DDBJ/ENA/GenBank under the accession NMPL00000000. The version described in this paper is version NMPL01000000. The PacBio and Illumina reads are available under the same BioProject. The TriticumD 1.0 contigs are available separately at ftp://ftp.ccb.jhu.edu/pub/data/Triticum_aestivum/Wheat_D_genome.

**Competing interests statement**. None of the authors have competing or conflicting interests.

## Acknowledgements

The authors wish to thank Jan Dvořák for helpful comments on the manuscript. We also acknowledge the support of the Johns Hopkins University large-scale computing center, MARCC, which provided invaluable computing resources. This work was supported in part by the U.S. National Science Foundation under grant IOS-1238231 and IOS-1444893, and by the U.S. National Institutes of Health under grant R01 HG006677.

